# The incidence of movement disorders increases with age and contrasts with subtle and limited neuroimaging abnormalities in argininosuccinic aciduria

**DOI:** 10.1101/2023.10.10.561631

**Authors:** Sonam Gurung, Saketh Karamched, Dany Perocheau, Kiran K. Seunarine, Tom Baldwin, Loukia Touramanidou, Claire Duff, Nour Elkhateeb, Karolina M. Stepien, Reena Sharma, Andrew Morris, Thomas Hartley, Laura Crowther, Stephanie Grunewald, Maureen Cleary, Helen Mundy, Anupam Chakrapani, Spyros Batzios, James Davison, Emma Footitt, Karin Tuschl, Robin Lachmann, Elaine Murphy, Saikat Santra, Mari-Liis Uudelepp, Mildrid Yeo, Patrick F. Finn, Alex Cavedon, Summar Siddiqui, Lisa Rice, Paolo G.V. Martini, Andrea Frassetto, Philippa B. Mills, Paul Gissen, Jonathan D. Clayden, Christopher A. Clark, Simon Eaton, Tammy L. Kalber, Julien Baruteau

## Abstract

Argininosuccinate lyase is integral to the urea cycle detoxifying neurotoxic ammonia and the nitric oxide biosynthesis cycle. Inherited argininosuccinate lyase deficiency causes argininosuccinic aciduria (ASA), a rare disease with hyperammonaemia and nitric oxide deficiency. Patients present with developmental delay, epilepsy and movement disorders, associated with nitric oxide-mediated downregulation of central catecholamine biosynthesis. A neurodegenerative phenotype has been proposed in ASA. To better characterise this neurodegenerative phenotype in ASA, we conducted a retrospective study in six paediatric and adult metabolic centres in the UK in 2022. We identified 60 patients and specifically looked for movement disorders-related symptoms: movement disorders such as ataxia, tremor and dystonia, hypotonia and abnormal behaviour. We analysed neuroimaging with diffusion tensor imaging (DTI) magnetic resonance imaging (MRI) in an ASA patient with movement disorders. We assessed conventional and DTI MRI alongside single photon emission computer tomography (SPECT) with dopamine analogue radionuclide ^123^I-ioflupane, in *Asl*-deficient mice treated by *hASL* mRNA with normalised ureagenesis. Movement disorders in ASA appears in the 2^nd^ and 3^rd^ decades of life, becoming more prevalent with ageing and independent from the age of onset of hyperammonaemia. Neuroimaging can show abnormal DTI features affecting both grey and white matter, preferentially basal ganglia. ASA mouse model with normalised ureagenesis did not recapitulate these DTI findings and showed normal ^123^I-ioflupane SPECT and cerebral dopamine metabolomics. Altogether these findings support the pathophysiology of a late-onset movement disorders with functional central catecholamine dysregulation but without or limited neurodegeneration of dopaminergic neurons, making these symptoms amenable to targeted therapy.

**Synopsis:** Movement disorders-related symptoms in ASA appear in the 2^nd^ and 3^rd^ decades of life, becoming more prevalent with age and shows abnormal neuroimaging features of basal ganglia in ASA patients, not recapitulated in ASA mice.

## Introduction

Argininosuccinic aciduria (ASA) (OMIM#207900) is a rare autosomal recessive metabolic disease, with a prevalence of 1/110,000 live birth and the second most common urea cycle disorder ^1^. Patients present with acute hyperammonaemia either in the neonatal period defined as early-onset phenotype, or later in life in late-onset presentation ^2^. Most patients develop a systemic phenotype with chronic neurological, hepatic and gastrointestinal problems, hypokalaemia and arterial hypertension ^3^. The neurological phenotype is variable with intellectual difficulties, attention deficit, epilepsy, behavioural changes and movement disorders ^3–5^. Although ammonaemia is neurotoxic and hyperammonaemia causes neurological sequelae, which can explain some of the symptoms observed in ASA chronic encephalopathy, these symptoms often occur despite satisfactory control of ammonia levels ^3,6^. Best-accepted therapy in ASA relies on ammonia control using protein-restricted diet, ammonia scavengers and arginine supplementation ^2,7^ with an increasing number of patients treated by liver transplantation ^8^. It was recently suggested that liver transplantation could have a sustained neurological benefit even in ASA patients with well-controlled ammonia levels ^9^.

ASA is caused by the deficiency of the cytosolic enzyme argininosuccinate lyase (ASL), the only mammalian enzyme enabling endogenous arginine synthesis ^10^. ASL breaks down argininosuccinic acid into arginine and fumarate, a biochemical reaction integral to the nitric oxide (NO) cycle, which enables NO synthesis from arginine, and the urea cycle, a liver-based pathway enabling nitrogen wasting through clearance of neurotoxic ammonia ^11^. The pathophysiology of ASA chronic encephalopathy is likely multifactorial with detrimental roles of hyperammonaemia, argininosuccinate toxicity ^12^, deficiency of arginine and downstream metabolites such as creatine ^1^, neuronal nitro-oxidative stress ^13^ and nitric oxide-mediated downregulation of central catecholamine biosynthesis suggesting a neurodegenerative phenotype ^14,15^.

Here, we present a national multicentre retrospective study assessing the phenotype of movement disorders in ASA, a well-recognised feature of ASA chronic encephalopathy. We provide a clinical description of neurodegeneration-related symptoms, movement disorders, hypotonia and abnormal behaviour, from a large cohort of patients, present neuroimaging data in humans and highlight the difficulties in modelling movement disorders in ASA mouse models. We show that the incidence of movement disorders in ASA increases with ageing, starting in adolescence and early adulthood. Abnormal diffusion tensor imaging (DTI) features can affect both the grey and white matter, preferentially close to basal ganglia. mRNA therapy-treated ASA mice with normalised ureagenesis did not recapitulate these DTI findings and showed normal single photon emission computer tomography (SPECT) with ^123^I-ioflupane, a dopamine analogue radionuclide. These findings support the pathophysiology of a movement disorders in ASA with functional central catecholamine dysregulation but without loss of dopaminergic neurons, making these symptoms amenable to targeted therapy.

## Material and Methods

### Patients

We conducted a retrospective study in six paediatric and adult tertiary metabolic centres in the UK. Epidemiological and clinical data of ASA patients with neurological disease were collected between July 2015 and June 2022 retrospectively from medical notes.

Movement disorders-related symptoms were considered when reported by a patient during medical history investigation or when observed during medical examination. The following symptoms were considered: (i) involuntary abnormal movements such as tremor or ataxia, excluding seizures, (ii) abnormal muscle tone, unusual and unexplained acute episodes of fatigue and chronic lethargy, (iii) early signs observed in neurodegenerative diseases such as sleep disturbances and abnormal behaviour.

Early-onset ASA was defined as hyperammonaemia occurring on/before 28 days of age, and late-onset ASA after 28 days of age in the patient or familial index case.

### Patient’s neuroimaging analysis

A brain MRI was performed in one ASA patient using the standard operating procedures of Great Ormond Street Hospital for Children with conventional T_1_-weighted and diffusion tensor imaging (DTI) sequences. Images were obtained using a Siemens Prisma 3T MRI scanner (Erlangen, Germany) and a 20-channel head coil. T1-weighted were acquired using an MPRAGE sequence with a TR of 2300ms, TE of 2.74ms, field of view of 256×256mm, 1mm slices and 256×256 matrix. The DTI sequence consists of a multiband SE-EPI sequence with 60 directions at b=1000 s mm^-2^ and 60 directions at b=2200 s mm^-2^, with 13 b=0 images interleaved throughout the sequence and an additional b=0 image with negated phase-encoding for distortion correction, TR of 3050ms, TE of 60ms, field of view of 220×220mm, a 110×110 matrix, 2mm slices with a 0.2mm slice gap and multiband factor 2. Basal ganglia volumes were derived from FreeSurfer^16^ (caudate, putamen and pallidum bilaterally) using an analysis of covariance with age, gender and total intracranial volume as covariates. Statistical analysis of FA and MD, averaged over each subcortical ROI was performed using an analysis of covariance with age and gender as covariates. Multiple comparisons correction was applied using false discovery rate (FDR).

### mRNA formulation

*hASL* and Luciferase (*Luc*) encoding mRNA encapsulated in Lipid Nanoparticles (LNPs) were provided by Moderna Therapeutics using their proprietary technology. Codon optimized mRNA encoding *hASL* was synthesized in vitro by T7 RNA polymerase-mediated transcription. The mRNA initiated with a cap, followed by a 5’ untranslated region (UTR), an open reading frame (ORF) encoding *hASL*, a 3’ UTR and a polyadenylated tail. Uridine was globally replaced with N1-methylpseudouridine, previously described ^17^. For in vivo intravenous delivery, LNP formulations were generated. Briefly, mRNA was mixed with lipids at a molar ratio of 3:1 (mRNA:lipid), previously described ^18^. mRNA-loaded nanoparticles were exchanged into final storage buffer and had particle sizes of 80 – 100 nm, >80% encapsulation of the mRNA by RiboGreen assay, and <10 EU/mL endotoxin levels.

### Animals

Animal procedures were approved by institutional ethical review and performed per UK home office licenses PP9223137 and PEFC6ABF1. *Asl^Neo/Neo^*mice (B6.129S7-*Asl^tm1Brle^*/J) were purchased from Jackson Laboratory (Bar Harbor, ME) and maintained on standard rodent chow (Harlan 2018, Teklab Diets, Madison, WI) with free access to water in a 12-hour light / 12 hours dark environment. WT littermates were used as controls and housed in the same cages. Genotyping was performed using DNA extracted from tail clips as previously described ^13^.

### Animal experimental design

*Asl^Neo/Neo^* animals were given systemic administration of *hASL* mRNA from birth then weekly until the age of 8 weeks at a dose of 1mg/kg or 2mg/kg for IV and IP injections, respectively. Untreated WT littermates were used as controls. All *Asl^Neo/Neo^* animals survived with normal growth and fur as previously described ^19^.

### Magnetic resonance imaging in mice

Images were acquired on a 9.4 Tesla Bruker imaging system (BioSpec 94/20 USR) with a horizontal bore and 440 mT/m gradient set with an outer/ inner diameter of 205 mm/116 mm, respectively (BioSpec B-GA 12S2), 86 mm volume coil, and a four-channel array receiver-surface coil (RAPI Biomedical GmbH). The brain regions of interest were first localized using a structural T2-TurboRARE sequence (fast-spin echo, Paravision 7.0). The olfactory bulbs were used as an anatomical landmark to maintain consistency in slice positioning between subjects and the slices covered the cortex and all subcortical structures up to the cerebellum. Imaging parameters for the T2-weighted imaging sequence were as follows: Repetition time (TR) = 4000ms, Time to Echo (TE) = 45ms, Field of view (FOV) = 21 x 16 mm^2^, data matrix 256 x 196, 14 x 600 µm coronal slices.

Diffusion weighted imaging (DWI) was performed using a 4-shot spin echo-planar imaging (EPI) sequence. Imaging parameters were: TR= 2500ms; FOV = 20 x 16 mm^2^; FOV = 20 x 16 mm^2^, data matrix 110 x 85; 14 x 600 µm coronal slices. To implement the multiple echo-time Neurite orientation dispersion and density imaging model (MTE-NODDI), the DWI images were acquired at three different echo times of 30, 45, and 60ms. At each echo time, MRI protocol consisted of two shells, detailed as follows:

1. Shell One: 30 directions, five b = 0 s/mm^2^ images and diffusion weighting of b = 2000 s/mm^2^
2. Shell Two: 15 directions, five b = 0 s/mm^2^ images and diffusion weighting of b = 700 s/mm^2^

with gradient duration and separation δ⁄Δ = 4.5⁄11 ms for all b-values and TE’s. The total acquisition time was approximately 120 minutes.

### Image processing and quantification of diffusion data

The effects of noise and imaging artifacts on the acquired diffusion data were reduced by applying a denoising method based on random matrix theory (MRtrix3), correction of B0 inhomogeneity and motion with TOPUP tool in FMRIB Software Library (FSL, University of Oxford, UK). DWI images were then co-registered to a reference b=0 image. Brain masks were created manually using ITK-SNAP software (www.itksnap.org) ^20^. Standard DTI measures of fractional anisotropy (FA) and mean diffusivity (MD) were generated from diffusion data obtained at TE = 30ms using dtifit in FSL, which fits a diffusion tensor model at each voxel of the data that has been pre-processed. (NODDI) parameters were estimated for each TE session separately with the NODDI MATLAB Toolbox, and the estimated parameters were aligned to the first TE session to extract TE-independent MTE-NODDI parameters.

For quantitative analysis, brain regions of interest (ROIs) were manually defined using ITK-SNAP. ROIs were drawn in WT and *hASL* LNP-mRNA treated Asl^Neo/Neo^ mice in the cortex, striatum for grey matter; corpus callosum, fimbria, and internal capsule for white matter; and hippocampus as a function region. All ROIs were subsequently exported to fitted diffusion maps and mean values for NDI, *f_en_*, ODI, FA, and MD were exported for quantitative analysis.

### SPECT imaging in mice

^123^I-ioflupane was obtained as a patient dose (185 MBq) from Curium Pharma UK Ltd. Mice were injected with approximately 15-20 MBq of ^123^I-ioflupane via tail vein injection. Mice were then anaesthetised using isoflurane anaesthesia (1.5-2% isoflurane in oxygen 1 L/min) and mouse head SPECT/CT scans were acquired 2 hours after injection using a NanoScan SPECT/CT system (Mediso, Hungary). CT images were acquired using a 55 kilo volt peak (kVp) X-ray source with 300 ms exposure time in helical mode, resulting in a scan time of approximately 3-4 minutes. SPECT images were obtained with a 4-head scanner with nine 1.4 mm pinhole apertures in helical scan mode using a time per view of 60 seconds resulting in a scan time of 40 minutes. Respiration was monitored throughout the scan and the body temperature was maintained by the heated bed. CT images were reconstructed in voxel size 124×124×124 μm using Mediso Nucline (Mediso, Hungary) software, whereas SPECT images were reconstructed in a 256 × 256 matrix using HiSPECT (ScivisGmbH, Bioscan/Mediso). Images were analysed using VivoQuant software (InViCro). 3D ROIs were drawn manually around to include the whole brain, a spherical region including the locus coeruleus, and basal ganglia. The percentage of injected dose/organ (%ID/organ) was calculated using decay-corrected ROI values.

### Dopamine metabolites

Dopamine, 3-O-methylDOPA (3-OMD), Homovanillic acid (HVA), 3,4-dihydroxyphenylacetic acid (DOPAC) and 5-hydroxyindolacetic acid (5-HIAA) were quantified using reverse-phase high performance liquid chromatography (HPLC) as previously described ^21^.

### Statistical analysis

Statistical analysis was performed using Prism 9.0 software (San Diego, CA, USA). Differences between groups were assessed using an two tailed unpaired *t* test with adjustment for multiple comparisons using false discovery rate (FDR). *p* values ≤ 0.05 were considered statistically significant. Correlation between continuous variables was assessed by Spearman’s rank correlation test.

## Results

### Demographic and Clinical characteristics

Sixty patients (32 males) were included with a median age of 12.7 years (range: 6 months-53 years). 34 patients (57%) had early-onset ASA. Diagnosis of ASA was obtained biochemically and was confirmed genetically in 25 patients (Supplementary Table 1). Three patients who died during the first month of life were excluded from the analysis.

### Clinical characteristics of movement disorders-related symptoms

This study included 17 (28%) ASA patients with neurodegeneration-related symptoms, movement disorders, hypotonia, abnormal behaviour, with a median age of 19 years (range: 4 years-53 years) and a sex ratio male/female of 10/7 (Table 1).

**Table 1.** Features of ASA patients with movement disorder-related symptoms. F Female, M Male, N No, NA Not available, Y Yes.

| Patient number | Form | Sex | Survival (Age at death) | Age at last follow-up (years) | Long term neurological phenotype |  |  |  |  |  |  |  | Genotype |
| --- | --- | --- | --- | --- | --- | --- | --- | --- | --- | --- | --- | --- | --- |
|  |  |  |  |  | Abnormal movements |  | Hypotonia |  | Behaviour |  | Intellectual disability | Epilepsy |  |
| 1 | Early | F | Alive | 20 | Y (Tremor, ataxia, dysdiadochokinesia) | 11y | Y (Fatigue) | 10y | N | / | Y | N | NA |
| 2 | Early | M | Alive | 4 | N | / | Y (Acute tiredness) | 1y | N | / | Y | N | c.1045-1057del; c.1045-1057del |
| 4 | Early | M | Alive | 12 | Y (Tremor) | 9y | Y (Myopathic face, fatigue) | 9y | Y (Agressivity, sleep disorder) | 17y | Y | Y | c.719-2A>G; c.857A>G |
| 5 | Early | M | Alive | 11 | N | / | Y (Acute tiredness) | 11y | N | / | Y | Y | c.437G>A; c.437G>A |
| 6 | Early | F | Died (12 years) | 13 | N | / | Y (Fatigue) | 13y | N | / | Y | N | NA |
| 8 | Early | M | Alive | 18 | N | / | Y (Fatigue) | 18y | N | / | Y | Y | c.1153C>T; c.1153C>T |
| 9 | Early | M | Alive | 25 | Y (Dystonia) | 25y | N | / | N | / | Y | Y | c.1153C>T; c.1153C>T |
| 14 | Early | F | Alive | 10 | Y (Ataxia) | 10y | N | / | N | / | Y | N | NA |
| 24 | Late | M | Died (53 years) | 53 | Y (Nystagmus, ataxia) | NA | Y (Swallowing difficulties) | NA | N | / | Y | Y | NA |
| 25 | Late | M | Alive | 24 | Y (Ataxia, tremor) | 24y | N | / | N | / | Y | Y | c.437G>A; c.446+1G>A |
| 27 | Late | F | Alive | 28 | N | / | N | / | Y (Agressivity) | 28y | Y | N | NA |
| 28 | Late | M | Alive | 16 | N | / | Y (Acute tiredness, fatigue, ptosis, myopathic face) | 12y | N | / | Y | Y | NA |
| 31 | Late | M | Alive | 19 | N |  | Y (Fatigue) | 15y | Y (Agressivity) | 16y | Y | N | NA |
| 33 | Late | F | Alive | 26 | Y (Ataxia) | 8y | N | / | N | / | Y | Y | c.348+1G>A; c.532G>A |
| 35 | Late | F | Alive | 22 | Y (Ataxia) | NA | N | / | N | / | Y | N | NA |
| 36 | Late | M | Alive | 20 | Y (Ataxia) | 9y | N | / | N | / | Y | N | NA |
| 47 | Early | F | Alive | 15 | N | / | N | / | Y (Agressivity, sleep disorder) | 10y | N | N | NA |

Movement disorders and hypotonia were the most frequent symptoms observed. Movement disorders were observed in 9 (15%) patients with a median age at onset of 10 years (range: 8 years-25 years) and a sex ratio male/female of 5/4 (Figure 1B). Tremor was observed in 3 patients and described as intention tremor, with high amplitude, rhythmic and oscillatory tremor during a purposeful movement. No resting tremor was observed. Patients with tremor reported a negative impact on their quality of life. No patient received tremor-targeting therapy. One patient presented with dystonia at the age of 25 year-old. Hypotonia-related symptoms were observed in 9 (15%) patients with a median age at onset of 11.5 years (range: 1 years-18 years) and a sex ratio male/female of 7/2 (Figure 1A). Behaviour changes were observed in 4 (7%) patients with a median age at onset of 16.5 years (range: 10 years-28 years) and a sex ratio male/female of 2/2 (Figure 1C). Some patients presented a combination of these symptoms (Figure 1D). After a free interval during childhood, the onset of hypotonia-related symptoms and behaviour changes was observed in the second decade, whilst movement disorders symptoms were recognised in early adulthood during the 3^rd^ decade (Figure 1E). These symptoms affected early- and late-onset patients similarly (figure 1F). The prevalence of these symptoms increased gradually with ageing with 100% of patients aged 25 to 29-year-old affected (Figure 1G). Patients aged >30 showed a lower prevalence of 33%. A correlation between prevalence of these symptoms and age was observed (Spearman’s correlation coefficient r^2^=0.22, *p*=0.34) (Figure 1H).

**Figure 1.**
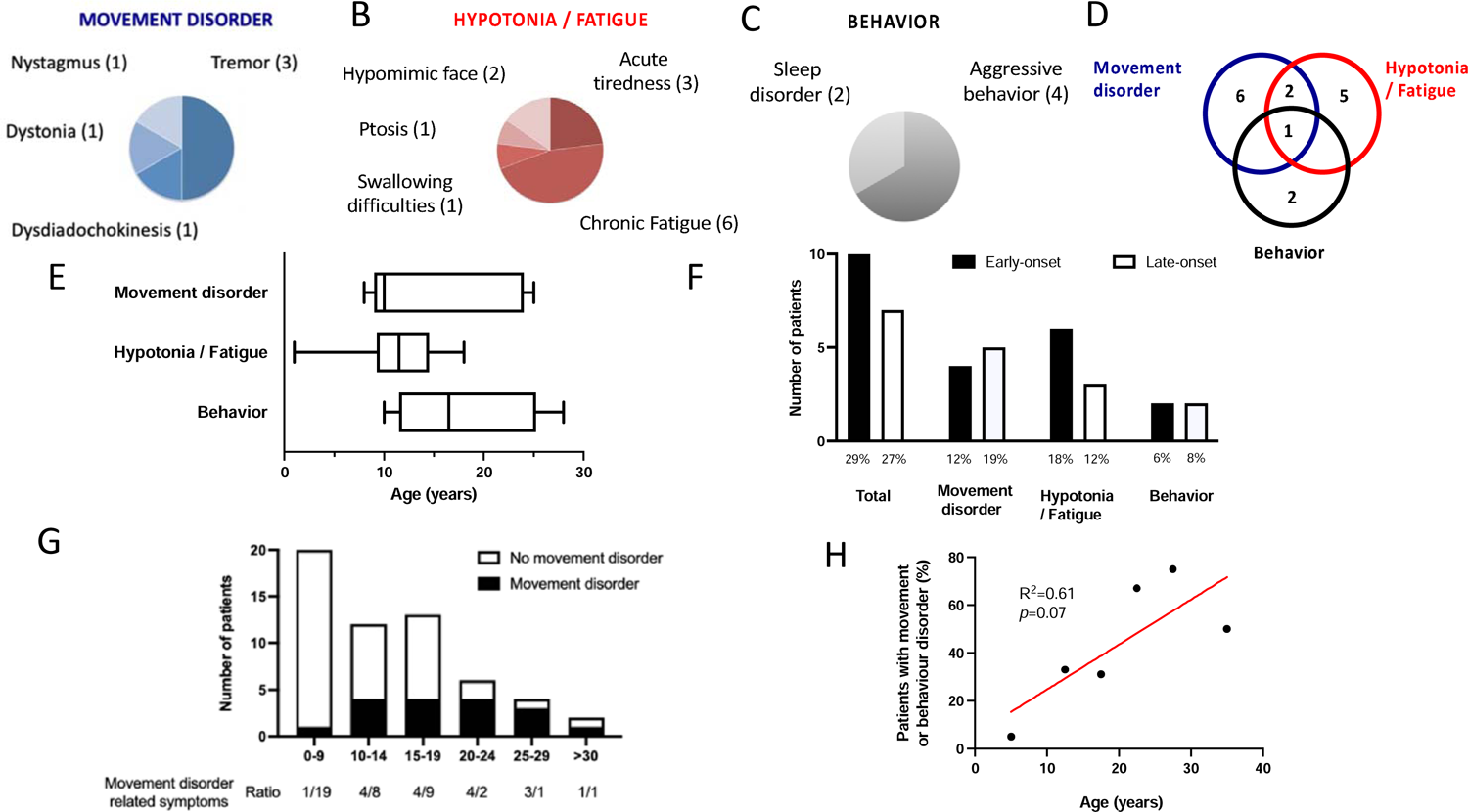
Movement, hypotonia and behaviour disorders in ASA patients. (**A**) ASA can present symptoms affecting (**A**) movement disorders, (**B**) hypotonia/fatigue and (**C**) behaviour. (**D**) These symptoms can be isolated or overlapping in one individual. (**E**) These symptoms are not present during the first decade and appear during the second and third decades. (**F**) The frequency of affected patients for these symptoms is similar between early- and late-onset ASA patients. (**G**) The ratio of ASA patients presenting with movement disorder, hypotonia/fatigue and/or behaviour is presented according to age range. (**H**) Simple linear regression between the percentage of individuals with ASA presenting with movement disorder, hypotonia/fatigue and/or behaviour against the age range of patients.

ASA patients presenting movement disorders-related symptoms did not have a higher risk of learning disability (*p*=1) and epilepsy (*p*=0.79), compared to ASA patients without movement disorders.

### ASA patient functional MRI showed involvement of basal ganglia, not identified in LNP-mRNA treated ASA mice

We investigated one neonatal-onset male ASA patient using brain MRI with conventional and diffusion tract imaging (DTI) sequences at the age of 16 years. The diagnosis was confirmed by identificastion of bi-allelic pathogenic mutations in the *ASL* gene (c.719-2A>G / c.857A>G/ p.Q286R) who was managed with protein-restricted diet, ammonia scavenger and arginine supplementation. He had recurrent hyperammonaemic decompensations during childhood and participated in a phase I/II clinical trial of cell therapy (NCT01765283), which did not show efficacy in improving ureagenesis and did not allow to relax the therapies based on standard of care. The patient had mild learning difficulties, with intention tremor, hypomimic face and occasional episodes of unexplained and self-resolving acute tiredness. This ASA patient had a brain MRI with conventional T_1_-weighted and diffusion tensor imaging (DTI) sequences at the age of 16 years to explore his motor disorder.

A global and detailed analysis with special interest on basal ganglia areas involved in movement disorders was performed and compared to age and sex-matched control cohort (N=21; 7 males; Age 16.7±1.7 years). The volume of the caudate, putamen and pallidum showed no difference between the ASA patient and the control cohort. The analysis of DTI MRI data focussed on the subcortical regions of interest showed a significantly higher MD for both left and right pallidum in the ASA patient, compared to controls (*p*=0.0006 and *p*=0.0022 respectively) as well as the left and right putamen (*p*=0.0037 and *p*=0.0041 respectively). No difference in FA was observed in the ASA patient compared to controls in any of the regions of interest. A tract-based spatial statistics (TBSS) analysis^22^ was performed to analyse the white matter microstructure. The patient has significantly lower FA than controls (p<0.05) bilaterally in the internal capsule, external capsule and cerebral peduncle, as well as the left anterior corona radiata and fornix (Figures 2A-C). No significant difference in MD for the white matter was detected.

**Figure 2.**
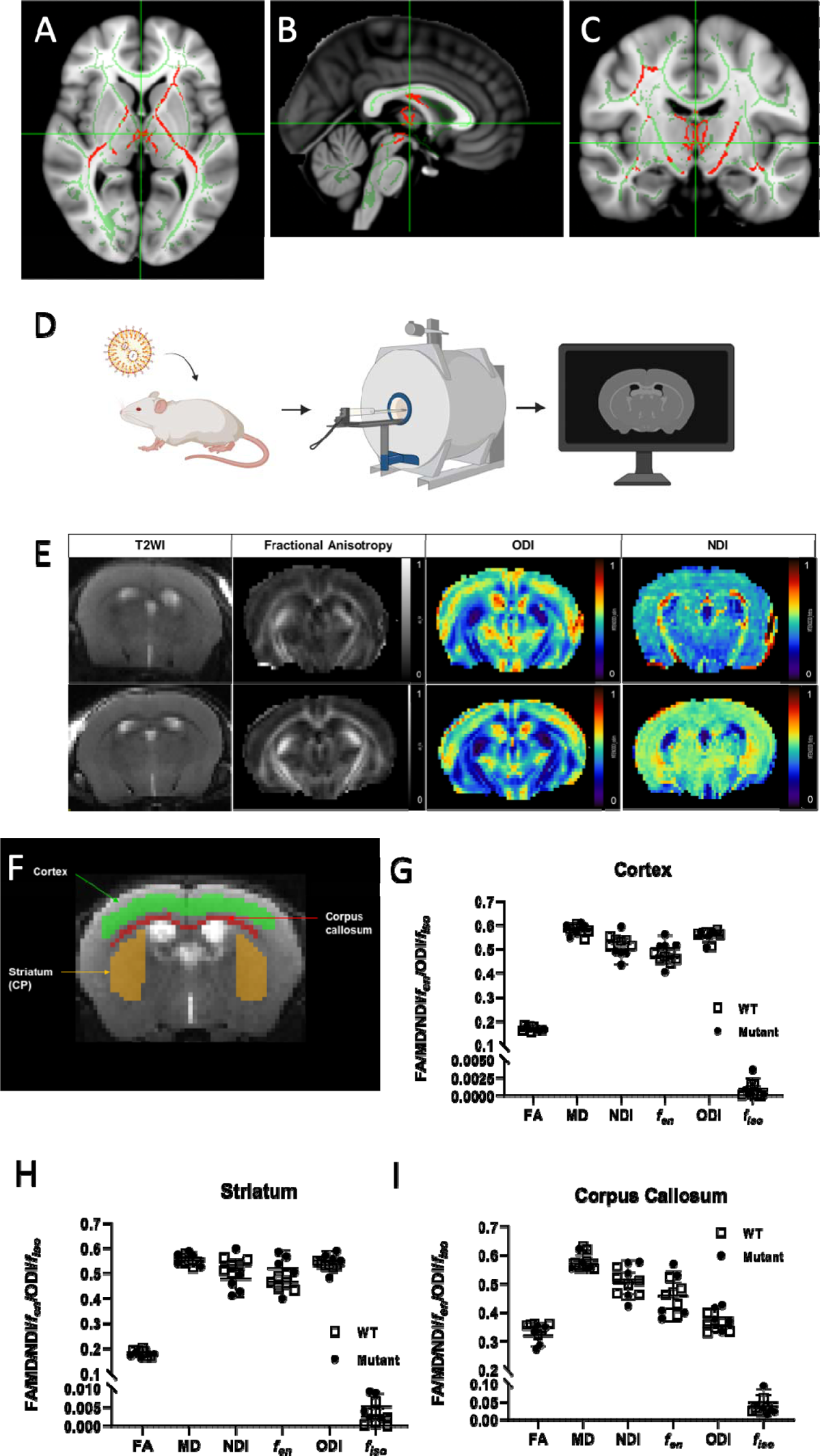
ASA patient functional MRI showed involvement of basal ganglia, not identified in LNP-mRNA treated ASA mice. (**A-C**) Decreased fraction of anisotropy in the internal capsule, fornix and white-matter tracts in the vicinity of basal ganglia in an ASA patient with (**A**) axial, (**B**) sagittal and (**C**) coronal sections. (**D**) Schematic describing the experimental study in *Asl^Neo/Neo^* mice treated systemically by *hASL* LNP-mRNA from birth until 7 weeks of age. The MRI was performed 3-4 days after the last injection of *hASL* LNP-mRNA, before post-processing and data analysis. (**E**) Representative MRI images of *hASL* LNP-mRNA treated *Asl^Neo/Neo^* mice with conventional T2, fractional anisotropy (FA), Orientation dispersion index (ODI) and Neurite dispersion index (NDI) sequences. (**F**) Manual definition of cerebral regions of interest with ITK-SNAP software. (**G-I**) Analysis of fMRI endpoints in various cerebral regions of interest (**G**) cortex, (**H**) striatum and (**I**) corpus callosum. Graph shows mean +SD.(**G-I**): Unpaired 2-tailed Student’s *t* test (False Discovery Rate corrected; p = 0.05; ns=not significant, n=5-6.

Modelling movement disorder-related symptoms is important to better understand the pathophysiology, especially which are the main cerebral structures affected, and test adequate therapies. To assess whether these abnormalities observed in an ASLD patient could be modelled in vivo, we used the *Asl^Neo/Neo^* mouse model, which is a recognised as a reliable model recapitulating the ASLD human phenotype. As we suspected that the ureagenesis defect is not the main cause of the motor phenotype in ASLD, we maintained this lethal *Asl^Neo/Neo^* mouse model by weekly systemic administration of *hASL* LNP-mRNA from birth until adulthood, when the animals underwent brain MRI with both structural and DWI sequences before downstream modelling and quantification (Figure 2D). The images obtained were of sufficient quality to reliably delineate the different brain structures and assess multiple parameters including volume on conventional T2-weighted images and standard DTI metrics such as fractional anisotropy (FA), mean diffusivity (MD). We also used the NODDI model to extract orientation dispersion index (ODI), extra-neurite volume fraction (f_en_), neurite dispersion index (NDI), and free-water volume fraction (isoVF) which describe underlying cerebral microstructure more accurately than DTI (Figure 2E). Different regions of interest were drawn (Figure 2F; Supplementary figure 1A) and analysed: frontal cortex (Figure 2G), striatum (Figure 2H), fimbria (Supplementary figure 1B) and hippocampus (Supplementary figure 1C) to represent grey matter; corpus callosum (Figure 2I), and internal capsule (Supplementary figure 1D) to assess white matter. In contrast to patient’s data (Figures 2A-C), no difference between WT and hASL LNP-mRNA treated Asl^Neo/Neo^ mice was observed (Figure 2G-I; Supplementary figure 1B-D).

### Dopamine metabolism in LNP-mRNA treated ASA mice

ASA patients develop disabling motor symptoms with age. The pathophysiology involves the NO-mediated downregulation of tyrosine hydroxylase and subsequent deficiency of central catecholamines ^15^. To date, different ASA mouse models have been described. The *Asl^Neo/Neo^*mouse is a hypomorphic mouse model with systemic disease. The *Asl^flox/flox^;TH Cre^+/-^* mouse is model of ASA with a dopaminergic neurone specific knock-out. Both models however exhibit motor abnormalities. Central catecholamine deficiency was demonstrated in *Asl^flox/flox^;TH Cre^+/-^*mice, but has not been assessed in *Asl^Neo/Neo^* mice. As *Asl^Neo/Neo^*mice recapitulates the human phenotype with systemic ASL deficiency affecting cerebral cell types, we aimed to model and assess the dopaminergic pathway *in vivo* using this model. We compared adult WT and *hASL* LNP-mRNA treated *Asl^Neo/Neo^* mice treated from birth with ^123^I-Ioflupane SPECT or DAT scan, which is routinely used in clinical settings to study the inherited ^23^ or acquired neurodegenerative ^24^ disorders affecting the dopamine synthesis or uptake pathway. Following intravenous administration of ^123I^-Ioflupane, no cerebral retention of the radiotracer was observed (Figures 3A, 3B). Analysis of ^123^I-Ioflupane retention index for the whole brain (Figure 3C), and uptake in locus coeruleus (Figure 3D) and basal ganglia (Figure 3E) did not show any difference.

**Figure 3.**
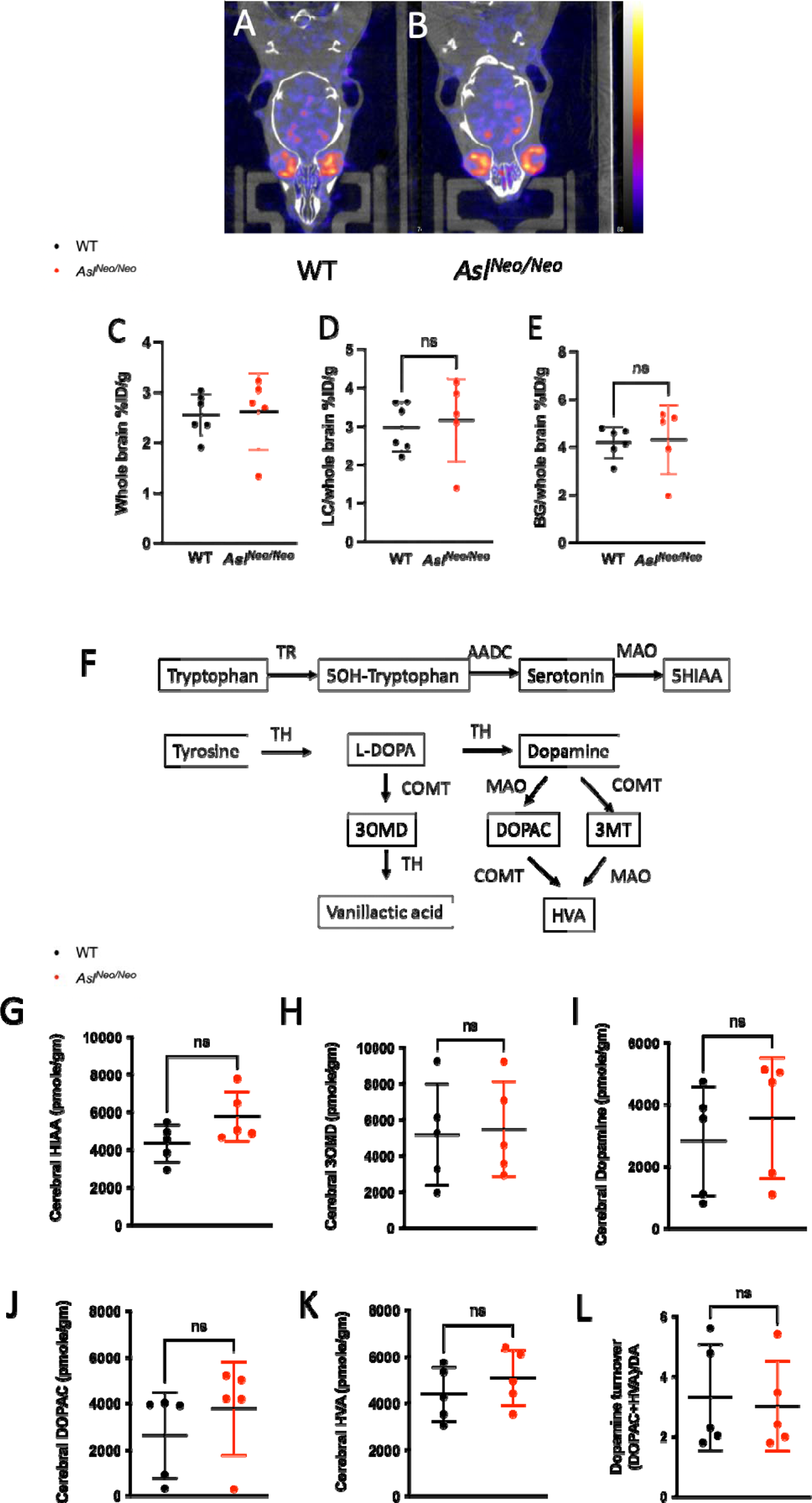
Dopamine metabolism in LNP-mRNA treated AS mice. (**A, B**) Representative pictures of DAT scan in WT and *hASL* LNP-mRNA treated *Asl^Neo/Neo^* mice. (**C-E**) Quantification of retention of ^123^I-ioflupane in (**C**) whole brain, (**D**) uptake in locus coereleus and (**E**) in basal ganglia. (**F**) Schematic representing the synthesis and catabolism of serotonin and dopamine. (**G-K**) Metabolomic data of serotonin and dopamine catabolism pathways in WT and *hASL* LNP-mRNA treated *Asl^Neo/Neo^* mice, (**G**) 5HIAA, (**H**) 3OMD, (**I**) dopamine, (**J**) DOPAC and (**K**) HVA. (**L**) Dopamine turnover estimated by (DOPAC+HVA)/Dopamine. Graph shows mean ±SD. (**C-E, G-L**): Unpaired 2-tailed Student’s *t* test; ns=not significant, n=5-6. DOPAC: 3,4-dihydroxyphenylacetic acid; 5HIAA: 5-hydroxyindolacetic acid; HVA: Homovanillic acid; 3OMD: 3-O-methylDOPA.

To assess whether serotonin and dopamine catabolism pathways (Figure 3F) were significantly disturbed in *Asl^Neo/Neo^* mice, cerebral dopamine-related metabolites were measured in 2-week-old untreated *Asl^Neo/Neo^*compared to WT mice and no difference was observed (Figures 3G-L). An increasing trend in cerebral 5-hydroxyindolacetic acid (5HIAA) levels was seen in *Asl^Neo/Neo^*mice compared to WT (*p*=0.09) (Figure 3G).

## Discussion

This study sheds light on the natural history of neurodegeneration-related symptoms, movement disorders, hypotonia, abnormal behaviour, which are common but poorly characterised features in ASA chronic encephalopathy ^3,15^. After a symptom-free period during childhood, neurodegeneration-related symptoms progressively develop during the 2^nd^ and 3^rd^ decades, affecting 29% of patients. Movement disorders including tremor and ataxia affected 15% of patients, which is comparable to 17% and 19% previously described by Baruteau *et al* in a British cohort ^3^ and Lerner *et al* in an international consortium with patients from North-America ^14^, respectively. These symptoms, especially intention tremor, are particularly disabling. Tremor is more characteristic of ASA when compared to other urea cycle defects ^14^. The main pathophysiological mechanism proposed is down-regulation of tyrosine hydroxylase (TH), the initial and rate-limiting step in the biosynthetic pathway of catecholamines including dopamine, noradrenaline, and adrenaline. TH activity is altered by two mechanisms: decreased ASL expression mediated by NO-mediated downregulation of TH transcriptional factor cyclic AMP (cAMP) response element-binding protein (CREB), and abnormal protein conformation caused by deficient nitrosylation ^15^. The pathogenic role of central catecholamine deficiency in ASA has been shown in abnormal stress response, epilepsy, memory and movement disorders. These symptoms can be partially rescued by NO donors ^14,15^.

Neuroimaging in ASA patients at baseline has shown various non-specific findings such as brain atrophy, focal infarcts, white matter and basal ganglia hyperintense signals, grey matter heterotopia and ulegyria and gliosis ^3,6^. In a significant number of cases (48%), neuroimaging with conventional MRI sequences is normal ^3^. MR spectroscopy can show reduced creatine or increased guanidino acetate in white matter and reduced N-acetylaspartate, suggesting reduced cellularity in basal ganglia ^3^. DTI enables a more accurate characterisation. In one patient with significant movement disorders reported here, brain MRI with conventional sequences appeared normal but DTI showed abnormal remodelling of grey and white matter affecting basal ganglia, especially globi pallidi, and white matter tracts in the vicinity of these structures. Pallidi are essential structures in movement control. Basal ganglia, especially thalami, are sensitive to hyperammonaemia ^25,26^ and abnormal signals in context of acute decompensations have previously been demonstrated ^27,28^. Although only one case is presented here, this may be characteristic of this disorder as abnormal basal ganglia findings are rare during follow-up of ASA patients with normal ammonia levels ^3,6^. Our case report highlights that DTI is a better tool than conventional MRI when assessing neuroimaging abnormalities in ASA patients with movement disorders. It is possible to speculate that previous hyperammonaemic decompensations in this patient are at least partially responsible for the DTI findings ^29^.

Further work is needed to provide definitive description of the nature and evolution of these abnormalities. Although this ASA patient was involved in a clinical trial with liver-directed cell therapy, no efficacy was observed, no improvement of ureagenesis was noted and the patient remained on the same combined therapy of diet and scavengers with similar dose.

We attempted to model the features observed in human neuroimaging in *Asl^Neo/Neo^* mice, which recapitulate the human disease phenotype and present motor and movement disorders. *Asl^Neo/Neo^* mice show systemic NO deficiency ^30^, especially in the brain ^13,31^, and is assumed to present with central catecholamine deficiency as observed in the knockout *Asl* model in dopaminergic neurons, *Asl^flox/flox^;TH Cre^+/-^* mouse ^14,15^. Conventional and DTI sequences did not show any significant differences between LNP-mRNA treated *Asl^Neo/Neo^*and WT mice. It could be that these abnormalities remain mild and are below the sensitivity of these neuroimaging tools. Additionally, liver-targeting mRNA therapy could as well partially alleviate the severity of the neurological disease, as suggested by in liver transplanted ASA patients ^9^. It is also possible that LNP-mRNA treated *Asl^Neo/Neo^* mice scanned at 8 weeks of life might have developed abnormal neuroimaging findings later in life. The absence of differences observed between LNP-mRNA treated *Asl^Neo/Neo^*and WT mice by cerebral ^123^I-ioflupane SPECT suggests the persistence of dopaminergic neurons with no neurodegeneration and supports the findings of Lerner *et al* with a functional reduction of central catecholamine synthesis, which responds to NO therapy ^14,15^. Another scenario could be a compensatory mechanism, with an increased number of DAT receptors to palliate the reduction of dopaminergic neurons as observed in Parkinson’s disease ^32^. Also, we cannot exclude the possibility that a SPECT imaging performed later in life could show a different result. The absence of differences in metabolomics analysis of dopamine synthesis and catabolism pathways between LNP-mRNA treated *Asl^Neo/Neo^* and WT mice suggests that TH downregulation remains compensated in this mouse model and does not affect the overall levels of central catecholamine metabolites. To better assess TH downregulation, measurement of dopamine metabolites and TH transcriptomics specifically in dopaminergic neurons rather than whole brain would provide a more accurate measurement. Overall, these findings bring hope for ASA patients with liver replacement therapy *i.e.* liver transplantation or gene therapy, as they support a therapeutic window, where movement disorders could be responsive to adequate therapy like NO donors, with no or limited damage of dopaminergic neurons.

This work has limitations due to the small number of patients identified with this rare disease, the methodology with retrospective analysis, the findings of DTI MRI reported in only one patient and limited evidence of TH downregulation in *Asl^Neo/Neo^*mice. Our findings require further prospective studies from larger cohorts of patients with urea cycle defects, which could be achieved via existing registries in Europe and in the USA, and warrant the inclusion of DTI sequences in neuroimaging of ASA patients ^33,34^. Additionally, to better decipher the complex neuropathophysiology of ASA, further characterisation of dopaminergic neurotransmission at the cellular or circuitry levels in *Asl^Neo/Neo^* mice would be required. Furthermore, developing surrogate models such as induced pluripotent stem cells derived neurons or 3-dimensional organoid cultures ^35,36^ or cultured *ex vivo* precision-cut organ slices ^37,38^ could become valuable alternatives to screen therapies able to treat TH downregulation, suspected to be the main driver of this movement disorders in ASA ^15^.

## Conclusion

In conclusion, neurodegeneration-related symptoms in ASA are a common and invalidating feature, which appear during adolescence and early adulthood. These symptoms are independent from the age of onset of hyperammonaemia DTI neuroimaging shows remodelling of basal ganglia, particularly globi pallidi, which provides an anatomical substratum for movement disorders. DTI neuroimaging, cerebral ^123^I-ioflupane SPECT and cerebral dopamine metabolomics in *Asl^Neo/Neo^* mice with restored ureagenesis failed to identify endpoints, which could be used for therapeutic testing, to target movement disorders.

## Authors’ contribution

SG and JB designed the project and wrote the manuscript. SG, SK, DP, LT, CD, TLK conducted or assisted with animal experiment and neuroimaging. KKS, CAK, JDC analysed patient’s neuroimaging. TB and SE performed the GCMS analysis of cerebral neurotransmitters. PBM and PG provided assistance with ethical agreement. NE, KMS, RS, AM, TH, LC, SG, MC, HM, AC, SB, JD, EF, KT, RL, EM, SS, MLU, MY, JB assisted with patients. PFF, AC, SS, LR, PGVM, AF provided LNP-mRNA formulation for animal experimental work. All authors reviewed and approved the final version of the manuscript.

## Competitive interest statement

JB is in receipt of research funding from Moderna Therapeutics. PFF, AC, SS, LR, PGVM and AF are employees of Moderna Therapeutics. Other authors have no competing financial conflict of interest to declare.

## Details of ethics approval

This research study was conducted retrospectively from medical notes. Participants’ data were recorded anonymously. Informed consent approved by the National Research Ethics Service Committee London-Bloomsbury (13/LO/0168) was obtained for participants and/or legal guardians for the following centres: Great Ormond Street Hospital for Children NHS Trust, National Hospital for Neurology and Neurosurgery, Evelina London Children’s Hospital, Salford Royal NHS Foundation Trust, and Manchester Centre for Genomic Medicine. Birmingham Children’s Hospital did not require consents from their institutional review board due to the collection of anonymous data.

## Consent for publication

Written consent for publication was obtained from the patient who underwent DTI neuroimaging.

## Data sharing statement

Database is available as supplementary material (Supplementary table 1).

## Approval from the Institutional Committee for Care and Use of Laboratory Animals

Animal procedures were approved by institutional ethical review and performed per UK home office licenses PP9223137 and PEFC6ABF1.

## Acknowledgments

We thank the patients for their participation in this study, and Mrs Megan Dorman and Mel McSweeney for their help in consenting patients.

