## Supplementary material for "The incidence of movement disorders increases with age and contrasts with subtle and limited neuroimaging abnormalities in argininosuccinic aciduria"

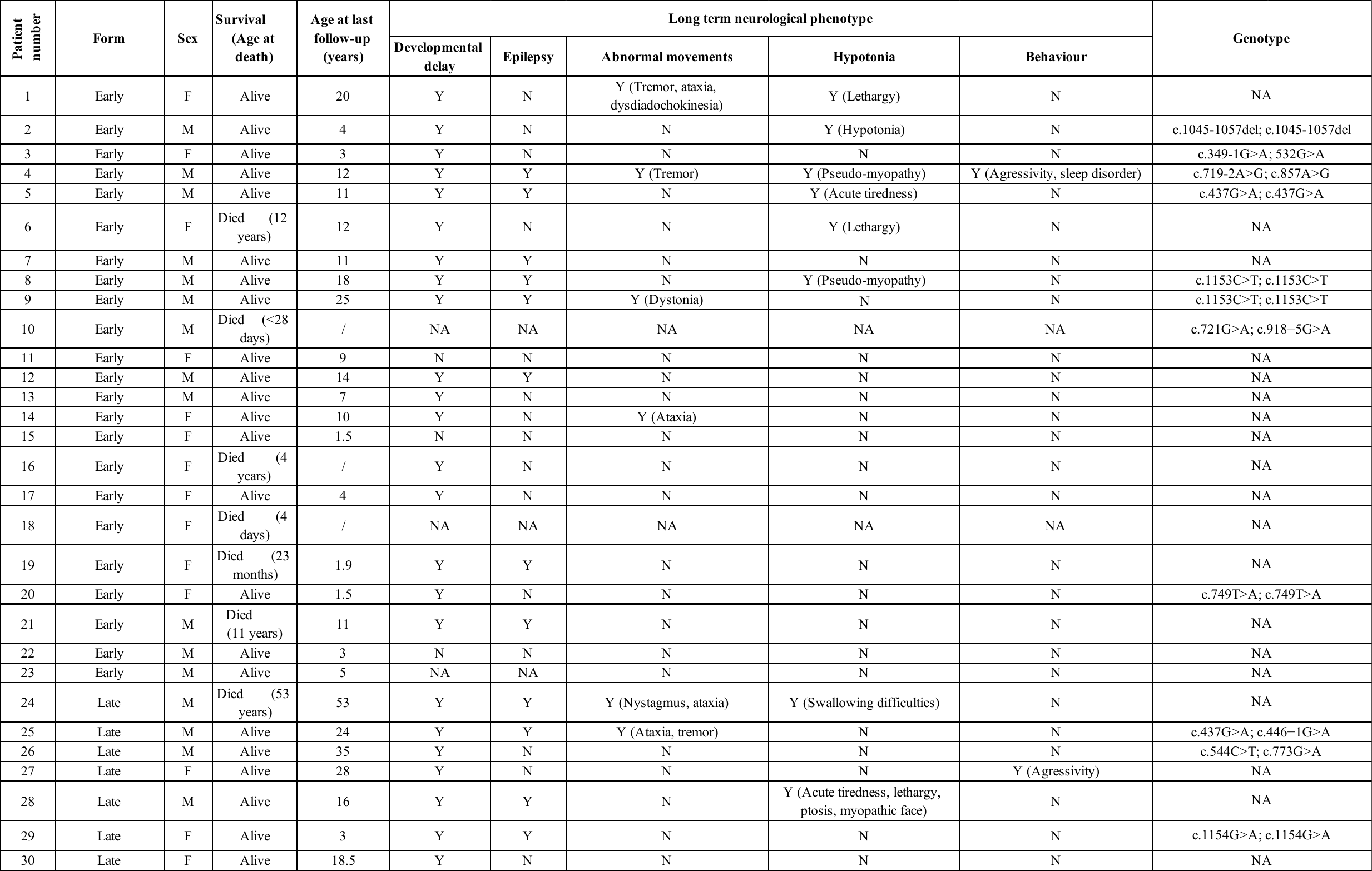

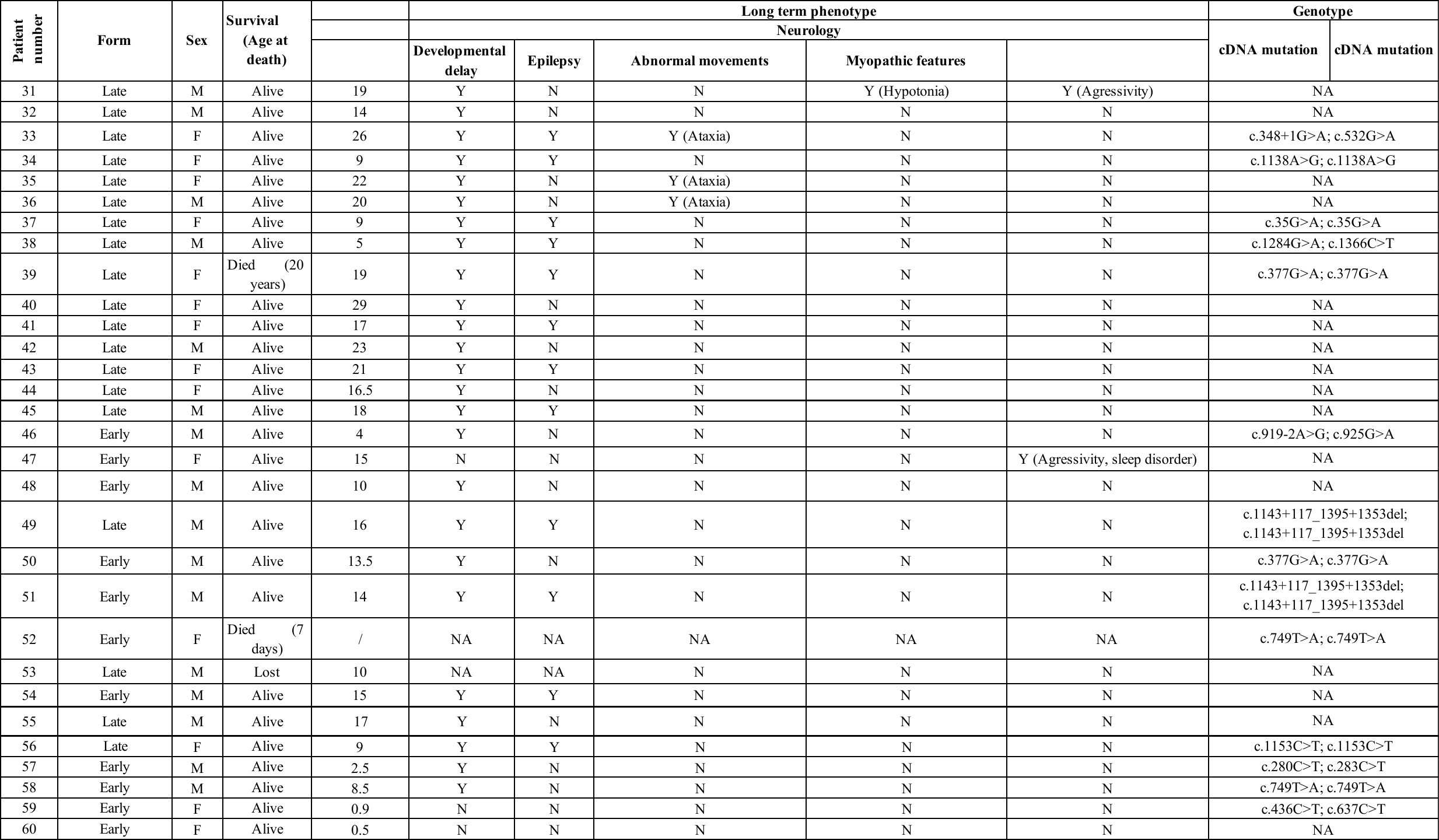

**Supplementary table 1. Features of ASA patients included in this study**

F Female, M Male, N No, NA Not available, Y Yes.

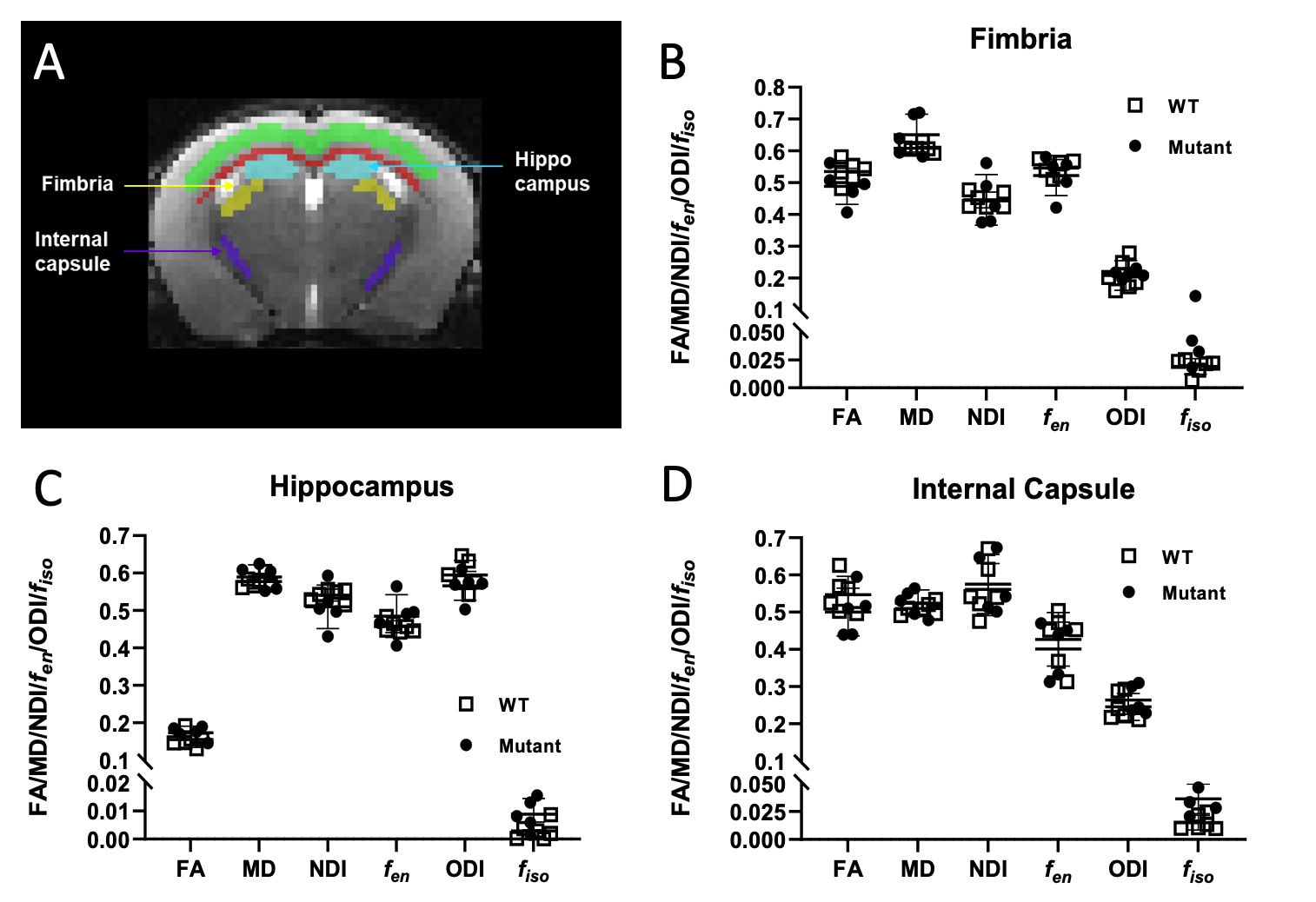

**Supplementary Figure 1. Functional MRI did not show involvement of regions of interest in LNP-mRNA treated ASA mice.** (**A**) Manual definition of cerebral regions of interest with ITK-SNAP software. (**B-D**) Analysis of fMRI endpoints in various cerebral regions of interest (**B**) fimbria, (**C**) internal capsule and (**D**) hippocampus. Graph shows mean $\pm$SD. (**G-I**): Unpaired 2-tailed Student’s *t* test (False Discovery Rate corrected; *p* = 0.05; ns=not significant, n=5-6.
